# gExcite - A start-to-end framework for single-cell gene expression, hashing, and antibody analysis

**DOI:** 10.1101/2022.05.23.490488

**Authors:** Linda Grob, Anne Bertolini, Matteo Carrara, Ulrike Menzel, Aizhan Tastanova, Christian Beisel, Mitchell P. Levesque, Daniel J. Stekhoven, Franziska Singer

## Abstract

**Summary:** Single-cell RNA sequencing (scRNA-seq) based gene expression analysis is now an established powerful technique to decipher tissues at a single-cell level. Recently, CITE-seq emerged as a multimodal single-cell technology capturing gene expression and surface protein information from the same single-cells, which allows unprecedented insights into disease mechanisms and heterogeneity, as well as immune cell profiling. Multiple single-cell profiling methods exist, but they are typically focussed on either gene expression or antibody analysis, not their combination. Moreover, existing software suites are not easily scalable to a multitude of samples. To this end, we designed gExcite, a start-to-end workflow that provides both gene expression and CITE-seq analysis, as well as hashing deconvolution. Embedded in the Snakemake workflow manager, gExcite facilitates reproducible and scalable analyses. We showcase the output of gExcite on a study of different dissociation protocols on PBMC samples.

**Availability:** gExcite is open source available on github at https://github.com/ETH-NEXUS/gExcite_pipeline The software is distributed under the GNU General Public License 3 (GPL3).

**Supplementary Information:** Supplementary information is available at the journal’s web site.

## Introduction

Over the past years, single-cell RNA sequencing (scRNA-seq) emerged as the next generation in high-throughput sequencing data analysis. Unlike bulk approaches, it offers insights on the gene expression on the single-cell level, allowing unprecedented insights in e.g. cell differentiation, immune compartment, and tumor heterogeneity (Kulkarni *et al*., 2019; Zhu *et al*., 2017). Recently, Cellular Indexing of Transcriptomes and Epitopes by Sequencing (CITE-seq) was introduced as a droplet-based single-cell profiling technology that enables the analysis of Antibody Derived Tags (ADTs) in addition to the gene expression (GEX) readout (Stoeckius *et al*., 2017). Taken together, CITE-seq can provide information on both the mRNA and surface protein level for the same cell. Moreover, hashing antibodies that tag ubiquitously expressed surface markers can be utilized for cell barcoding, allowing the multiplexing of samples. With this, a higher throughput of cells can be analyzed together, thereby decreasing the overall per-sample processing costs (Stoeckius *et al*., 2018). Consequently, the bioinformatics workflow needs to provide a demultiplexing step before entering the analyses of individual samples.

A variety of methods exists for the analysis of scRNA-seq data, including the widely used Seurat (Hao *et al*., 2021) and Scanpy (Wolf *et al*., 2018) suites, as well as pipelines such as scAmpi (Bertolini *et al*., 2021) and CReSCENT (Mohanraj *et al*., 2020). In addition, methods for either CITE-seq analysis or hashing demultiplexing are available, e.g. Seurat (Hao *et al*., 2021).

However, despite the availability of tools for the individual analysis, a workflow that combines the analysis of hashing, GEX and ADT data would greatly aid and simplify the integrated analysis for each cell. To this end, we implemented gExcite (pipeline for Gene EXpression and CITE-seq analysis), a workflow that facilitates hashing demultiplexing, and individual as well as combined ADT and GEX analysis from raw reads up until readily interpretable output such as cluster and cell type identification with combined information on gene and protein expression. Moreover, gExcite offers a customizable template for GEX and ADT-based differential expression analysis. All steps of the gExcite workflow are embedded into the Snakemake workflow manager (Mölder *et al*., 2021), following the latest Snakemake best practices (https://snakemake.readthedocs.io/en/stable/snakefiles/best_practices.html), and taking advantage of its framework for using conda (https://anaconda.com/) environments. Thereby, gExcite presents a scalable and reproducible pipeline for combined hashing, GEX, and ADT analysis.

## Workflow

The workflow implemented in gExcite can be conceptually divided as shown in Figure 1.

**Fig 1:**
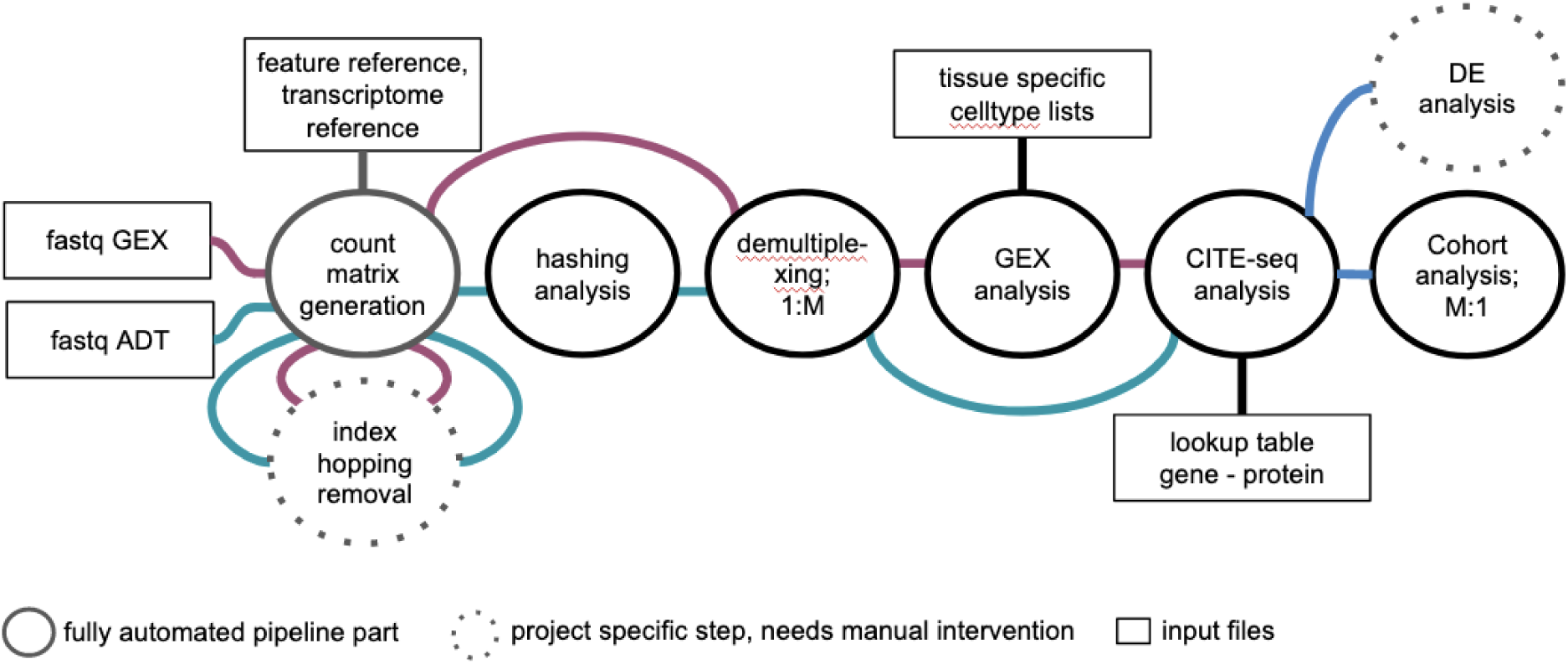
Overview of the workflow implemented in gExcite starting from raw fastq GEX and ADT files up until a combined analysis of GEX and ADT.

### Read mapping

Raw reads are mapped to a reference genome using 10x Genomics Cell Ranger software (Zheng *et al*., 2017). GEX and ADT libraries are processed independently to infer read counts per gene per cell, or read counts per antibody per cell, respectively. Required input files for this step include the fastq files as well as the transcriptome and feature reference.

If required by the experimental setup (e.g., sequencing on the NovaSeq without dual indexing), gExcite provides functionality to perform index-hopping removal using DropletUtils’ (Griffiths *et al*., 2018) ‘swappedDrops’ function. Based on the preliminary count matrix, index-hopping removal is performed resulting in a clean count matrix that can be further processed within the gExcite workflow.

### Hashing

Cell hashing allows for multiplexing, facilitates doublet detection and allows superloading of cells in single-cell transcriptomics experiments (Stoeckius *et al*., 2018).

In gExcite, the Seurat framework (Hao *et al*., 2021) is utilized to demultiplex samples after cell hashing and remove doublets as well as negatives (i.e., cells that cannot be assigned to any hashtag). First, CITE-seq-Count (Mimitou *et al*., 2018) is applied to count hashtags per cell. Subsequently, normalization and demultiplexing are performed using Seurat. For the normalization step itself different options are available: normalization across hashtags (features) or cells. In addition, gExcite offers a combined normalization option where the counts are normalized across cells and hashtags independently, and the resulting hashtag assignments are compared. Flexible settings are available on how to proceed with disagreeing hashtag assignments (e.g., cells with a disagreeing hashtag assignment are labeled as doublets). This combinational approach strengthens the confidence into the final hashtag assignment that separates the cells of different experimental settings. Refer to Supplementary Materials S1 for further details on the hashing analysis and for an illustration of the effects of different normalization options.

### GEX analysis

gExcite includes the scRNA-seq analysis pipeline scAmpi as a Snakemake module to analyze GEX data, including among other analyses cell type assignment, clustering, and pathway enrichment analysis. Importantly, the quality control functionality of scAmpi is used to further filter the GEX data and to remove contamination, such as likely empty droplets or ambient RNA (Bertolini *et al*., 2021).

### CITE-seq analysis

In the previous GEX analysis step the GEX data has been filtered as part of the quality control. Only cells present in both GEX and ADT data are preserved for downstream analysis. Thus, the ADT data is filtered based on the results of the GEX quality control.

The raw counts of the demultiplexed and filtered samples are log-transformed. gExcite offers functionality to analyze and visualize antibody expression data on its own as well as in combination with the corresponding gene expression (Supplementary Figure S2A, panel C and D). The link between antibodies and corresponding genes is provided by the user with a gene to protein dictionary. Gene and antibody expression counts can be simultaneously visualized in a Uniform Manifold Approximation and Projection (UMAP) plot (Supplementary Figure S2A, panel A). The UMAP embedding is computed either based on gene expression, antibody expression, or the combination thereof (Supplementary Figure S2B). This allows the assessment of cell similarity based on the different data types. BremSC (Wang *et al*., 2020) is utilized to provide a combined gene and antibody based clustering (Supplementary Figure S2A, panel B). Note that for CITE-seq analysis raw counts can contain background noise from ambient antibodies and nonspecific antibody binding. gExcite allows setting an experiment-specific threshold per antibody to accommodate this noise (Supplementary Figure S2A, panel D and F). Cells exceeding the threshold are labeled as positive for the respective antibody. To aid manual threshold definition and to ease interpretation of antibody expression, gExcite provides cell type-expression ridge plot visualization (Supplementary Figure S2A, panel E and F).

### Differential expression analysis and cohort-level analysis

We provide functionality for differential expression (DE) analysis on GEX and ADT data. Output from the gExcite workflow, in the form of either SingleCellExperiment (Amezquita *et al*., 2020) or Seurat R data objects, can be directly provided to the differential expression scripts to retrieve the list of genes or antibodies that pass user-defined expression and significance thresholds. The user has the freedom to choose between two DE analysis approaches: “pseudo-bulk” using DESeq2 (Love *et al*., 2014) or Seurat FindMarkers (Hao *et al*., 2021).

Regardless of the procedure, the output comprises the full list of tested genes with the test results, quality control plots (Supplementary Figure S3, panel A and panel B), expression barplots for the top 40 differentially expressed genes (Supplementary Figure S3, panel C), expression heatmap of a user-defined number of top differentially expressed genes with sample and contrast annotation (Supplementary Figure S3, panel D). More information can be found in Supplementary materials S3.

In addition, gExcite offers functionality to collectively analyze samples multiplexed within one experiment. Scripts are provided (refer to the github repository, folder “scripts/aggregation_DE”) that facilitate pooling individual samples, adding available metadata, and performing basic plotting and clustering. As such analyses are typically very project-specific, these scripts only provide a template that needs to be adapted according to the research question at hand.

## Conclusion and outlook

The proposed workflow offers comprehensive functionality for the automated analysis of both GEX and ADT, as well as hashed single-cell data. It facilitates an easy yet in-depth quality control of the analyzed samples as well as supports the interpretation of single-cell experiments. Key aspects are its flexibility, ease-of-use, and scalability, which allows the reproducible application also to large-scale data sets.

## Acknowledgements

The authors thank Michael Prummer and Lourdes Rosano for their support in GEX analysis features. CB’s lab was supported by the ETH domain Personalized Health and Related Technologies (PHRT-510).

## Author contributions

*LG+AB: pipeline design and implementation; manuscript*

*MC: implementation differential gene expression analysis; manuscript*

*LR + MP: pipeline part implementation*

*UM+ AT+ML+CB: showcase data generation and CITE-seq feedback*

*DS: manuscript review*

*FS: pipeline design; manuscript*

