## Supplementary material for "gExcite - A start-to-end framework for single-cell gene expression, hashing, and antibody analysis": Supplemetary Material S1

### S1: Hashing analysis

If cell hashing is performed, each cell barcode has to be assigned to one of the input samples (demultiplexing) before further downstream analyses of the individual samples. In the preprocessing for the demultiplexing, the ADT counts have to be normalized.

The count normalization with the Seurat function `NormalizeData()`, can be performed for each feature independently (with normalization.method = "CLR", margin = 1), or for each cell independently (with normalization.method = "CLR", margin = 2). The resulting cell barcode assignments to the hashing tags are often very similar, but can increasingly differ if e.g. one hashtag performs inferior to the others, or the experiment was sequenced at shallow depth. Therefore, gExcite provides options (config section: analyse_hashing) to combine the two normalization approaches. E.g., both normalization approaches can be performed independently and afterwards the respective cell barcode assignments to the sample hashtags are compared. If the assignment is identical between normalization approaches, it is kept as the final classification (e.g. "Doublet" in cell-based and tag-based normalization is kept as "Doublet" in the combination). If the assignments disagree, the user can fine-tune the final decision based on specific parameter settings, e.g. with the "save_negatives" option to either strictly remove negatives or prefer the singlet hashtag assignment. Table S1 gives an overview of how the different options would classify a cell in case the two normalization schemes differ.

| Normalization per feature | Normalization per cell | final classification using `--save_negatives FALSE` | final classification using `--save_negatives TRUE` | final classification using`--normalisation_downstream 1` | final classification using `--normalisation_downstream 2` |
|---|---|---|---|---|---|
| Negative | doublet | | | Negative | doublet |
| Negative | tag_A | Negative | tag_A | | |
| Doublet | tag_A | Doublet | Doublet | Doublet | Doublet |
| tag_A | tag_B | Doublet | Doublet | Doublet | Doublet |

***Table S1: Overview of hashtag assignments combining two normalization methods in the hashtag demultiplexing.*** *"Final classification" refers to the different parameter combinations of combining the two normalization approaches.*

A variety of quality control plots are generated for the demultiplexing step, to assist in identifying issues with the sample quality (see Figure S1). The plots allow the comparison of the performance of each hashtag and to check the underlying hashtag oligo (HTO) counts of the cells.

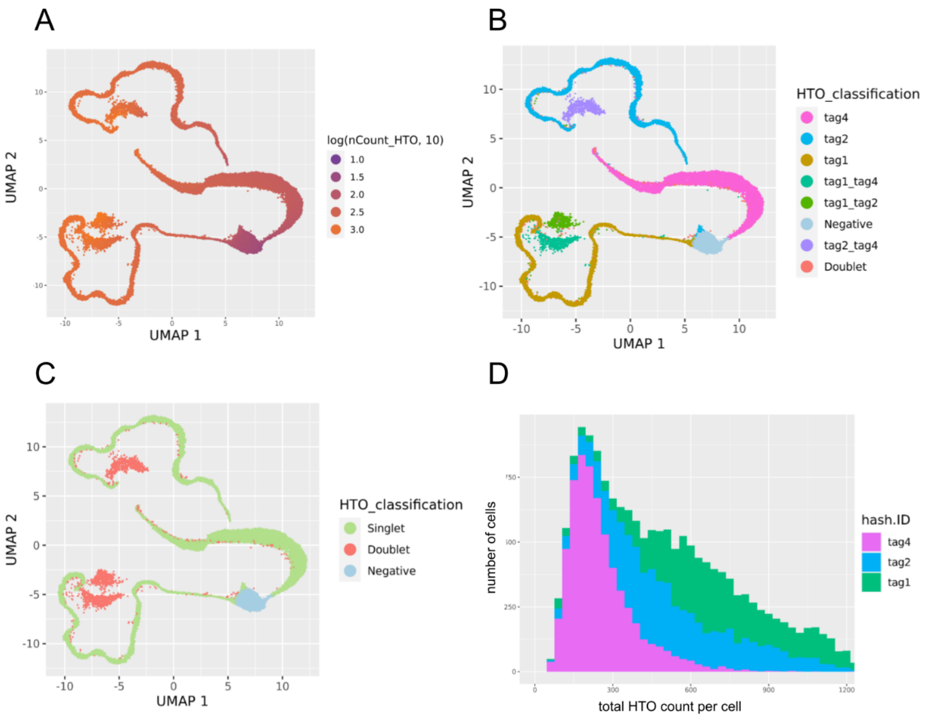


***Figure S1: QC plots of the hashtag demultiplexing step**.*

*In (A), the UMAP visualization is calculated based on raw counts, and the coloring shows the total ADT log count of each cell. In (B), the coloring shows the final hashtag classification assigned to each cell. (C) shows the UMAP colored by the general hashtag category assigned to each cell. (D) shows the histogram of total raw counts per cell, with bars coloured by assigned hashtag.*
