## Supplementary material for "gExcite - A start-to-end framework for single-cell gene expression, hashing, and antibody analysis": Supplemetary Material S2

### S2: CITE-seq Analysis

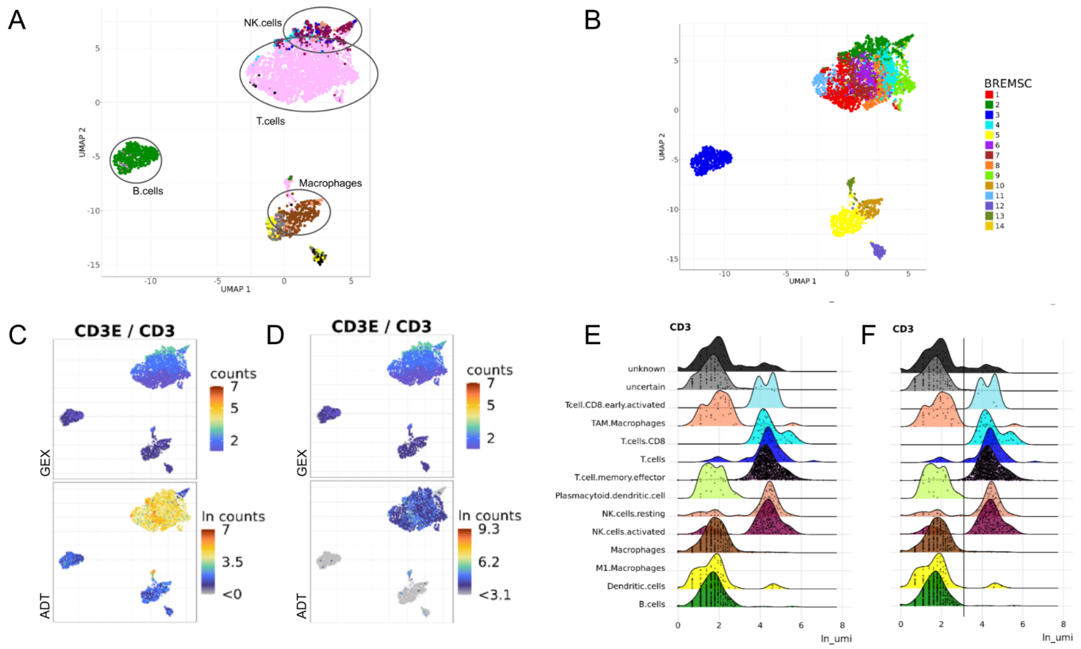


***Figure S2A: Showcase example of CITE-seq analysis.***

*In (A), the UMAP based on the combination of gene and antibody expression counts is coloured by the assigned cell type based on scAmpi's cell typing algorithm. In (B), the UMAP shows the clustering assigned by BREMSc, a clustering algorithm that includes scRNA and ADT information. (C/D) shows the UMAP colored by the counts of an example gene (top) and protein (bottom) combination. In the (C) bottom UMAP the normalized log counts of the ADT are shown as given out by TOOL, whereas in (D) the chosen/deduced/manually set threshold for background noise (shown in F) was subtracted from the protein counts. The ridge plots in E and F show the density of the ADT counts split up by cell type. In (F), the chosen threshold for this ADT molecule is marked as a horizontal line.*

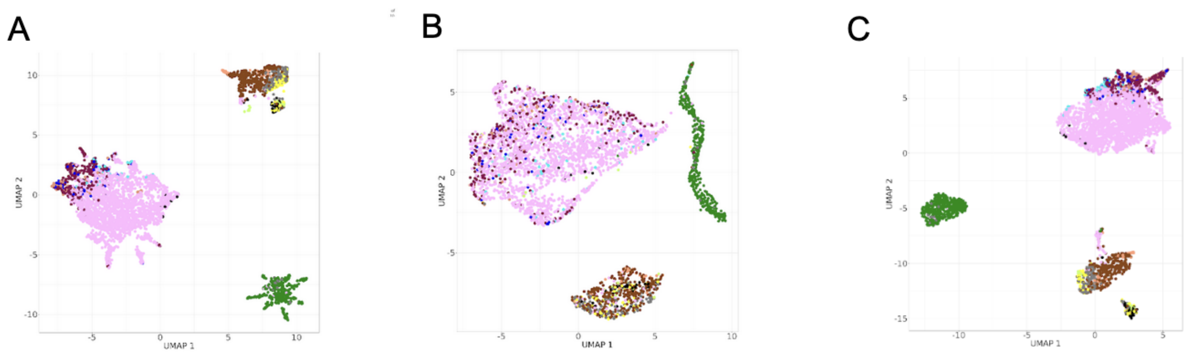


***Figure S2B: Showcase example of different UMAP embeddings.***

*The UMAP embeddings are computed either based on gene expression (A), antibody expression (B), or the combination (C) thereof.*
