## Supplementary material for "gExcite - A start-to-end framework for single-cell gene expression, hashing, and antibody analysis": Supplemetary Material S3

### S3: Differential expression

We provide two conceptually different approaches for differential expression analysis of single-cell data, each with different advantages and disadvantages: “pseudo-bulk” and Seurat FindMarkers.

Briefly, the “pseudo-bulk” approach relies on aggregating data from multiple samples by cell type in order to simulate the expression distribution of bulk RNA-sequencing. This ensures that the data have characteristics and a distribution that match the assumptions of a standard DESeq2 workflow. In contrast, Seurat FindMarkers is specifically designed for single-cell data and is based on the non-parametric Wilcoxon rank sum test.

Evidence shows that pseudo-bulk approaches tend to outperform single-cell-specific methods with a better false positive rate for highly-expressed genes (Squair *et al.*, 2021) Literature is, however, lacking a similar comparison on ADT data, which is differing from GEX data in the way that it has a much smaller number of features, but with less sparse counts.

Regardless of the chosen method, the scripts analyze pairwise contrasts with or without covariates. The output of the analysis provides:

- The full list of genes or antibodies with their associated differential expression statistics and metrics, along with a custom classification based on the user-defined fold-change and P-Value thresholds (default thresholds: fold-change = 1.5, P-Value = 0.05)
- A P-Value distribution barplot and a volcano plot to be used as quality controls
- Expression boxplot panel of the top 40 genes or antibodies, ranked by P-value
- Expression heatmap of the involved genes or antibodies

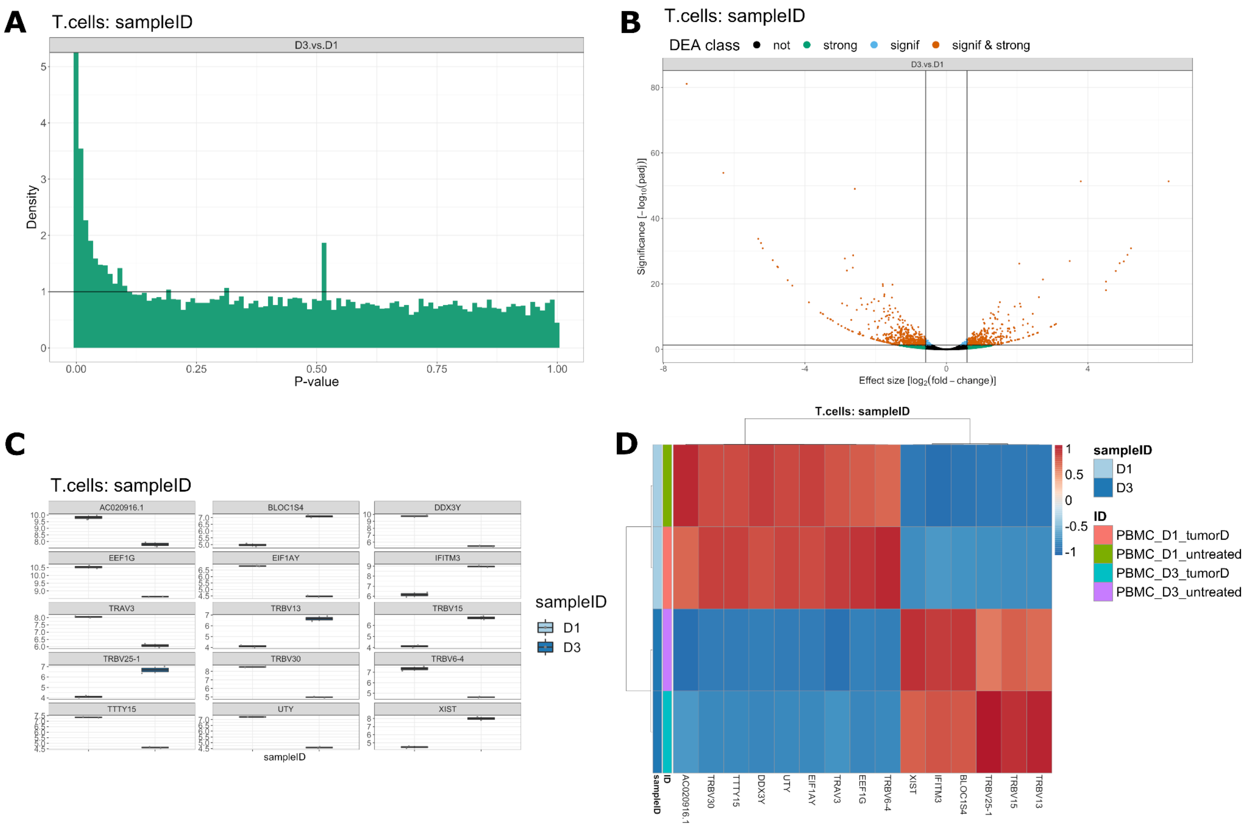


***Figure S3: Showcase examples of differential expression analysis output.***

*The differential expression analysis is applied independently on both GEX and ADT data. The output provides figures for quality control as well as figures to aid interpretation of the DE results. In (A), the barplot shows the distribution of P-values for the contrast. In (B), the Volcano plot shows the relationship between the effect size (log2 fold change) and the P-values for each tested gene. Genes are color-coded based on the user-defined thresholds for both metrics. In (C), the expression boxplots of up to 40 top significant genes compare the expression levels available for the two variables in the pairwise contrast. In (D), the expression heatmap shows the expression of top genes in all samples used to compute the differential expression analysis.*
